# IFN-I signaling in type 2 conventional dendritic cells supports T_H_2 and T follicular helper differentiation after allergen immunization

**DOI:** 10.1101/2024.09.10.612251

**Authors:** Greta R Webb, Kerry L Hilligan, Sam I Old, Shiau-Choot Tang, Olivier Lamiable, Franca Ronchese

## Abstract

Type 2 dendritic cells (DC2s) are essential for T_H_2 differentiation, but the signaling pathways involved in allergen sensing, DC activation and instruction of CD4+ T cell priming remain unclear. Previous transcriptomic analyses demonstrated a type-I interferon (IFN-I) signature in skin cDC2s following immunization with non-viable larvae of *Nippostrongylus brasiliensis* (*Nb*), house dust mite (HDM), and *Schistosoma* egg antigen (SEA). Blocking IFN-I signaling with anti-IFNAR1 (aIFNAR1) led to reduced T_H_2 cytokine responses to these antigens, however, the phenotype of cytokine-producing CD4+ T cells was not further defined. Here we show that conditional loss of IFNAR1 signaling in CD11c+ DCs significantly impaired effector T_H_2 and TFH CD4+ T cell responses to *Nb*. *In vivo* proliferation experiments demonstrated reduced numbers of highly divided CD4+ T cells in IFNAR1^ΔCD11c^ mice compared to IFNAR1^WT^, with the highly divided population comprising both T_H_2 and TFH. Characterization of the cDC2 compartment by flow cytometry and bulk RNAseq demonstrated lower numbers of *Nb*+ cDC2s in the skin-draining LN and a reduced expression of *Il15* and *Il15Ra* in IFNAR1^ΔCD11c^ mice compared to IFNAR1^WT^, while expression of costimulatory molecules including CD80, CD86, *Cd40* and *Pdcd1lg2* (PD-L2) was not impaired. Therefore, IFN-I conditioning of skin cDC2s is necessary for their effective priming of CD4+ T cell responses to allergens, providing evidence for a role of tissue cytokines in driving cDC2 activation in a T_H_2 context.

## Introduction

T_H_2 differentiation is driven by migratory conventional type-2 dendritic cells (cDC2s), which carry allergens from the site of exposure to the draining lymph node (dLN) and present them to naïve CD4+ T cells (1, 2). In the skin and skin-draining LN, migratory cDC2s can be divided into CD11b-hi and CD11b-low subsets. While both subsets of cDC2s can take up allergens (2, 3), the KLF4-dependent CD11b-low cDC2s have been shown to be essential for T_H_2 responses to *Schistosoma* egg antigen (SEA) (4), *Nippostrongylus brasiliensis* (*Nb*), and house dust mite (HDM) (3).

As allergens are mostly unable to induce direct DC activation, inflammatory cytokine production in peripheral tissues is thought to play an important role in cDC2 conditioning. For instance, Thymic Stromal Lymphopoietin (TSLP) was shown to condition DCs, and especially CD11b-low cDC2s, to the induction of T_H_2 differentiation to contact sensitizers (5, 6) while IL-33 and IL-13 can condition cDC2s to promote allergic and anti-helminth responses in the lung and gut (7, 8, 9, 10, 11). We previously demonstrated a robust type-I interferon (IFN-I) signature in skin cDC2s from mice immunized intradermally with non-viable *Nb* larvae, suggesting a role in cDC2 conditioning for T_H_2 differentiation (12). Antibody-dependent blockade of the IFN-I receptor IFNAR1 prevented the upregulation of IFN-I response markers on DCs and significantly impaired the IL-4 response to *Nb* (12). Similarly, *Schistosoma* egg antigen-loaded IFNAR1-KO bone-marrow derived DCs were unable to promote the differentiation of IL-4+ CD4+ T cells after *in vivo* transfer (13).

IFN-I signaling through the IFNAR1 and IFNAR2 receptor complex leads to the activation of multiple signaling pathways including IRF1 (14), IRF3 (15), IRF7 (16) and IRF8 (17), NFκB (18), STAT1 (19), STAT2 and STAT4 (20). IFN-I plays a fundamental role in cDC function during a variety of immune responses, and can increase proteasomal activity (21, 22), antigen presentation to CD8+ T cells by both cDC1s and cDC2s (23, 24), cDC2 migration, costimulation (13, 25), and metabolic activation (26, 27). IFN-I was also necessary for the development of CD64+ “inflammatory” cDC2s, which display enhanced CD4+ T cell priming capabilities following acute viral infection in the lung and are also observed following intradermal immunization with HDM and *Nb* (17). IFN-I signaling in cDC2s has been shown to be crucial for the differentiation of TFH in mesenteric LN following OVA + LPS challenge (28), as well as in the skin-draining LN following viral infection (29). Despite its demonstrated role in cDC biology and a range of CD4+ T cell responses, the mechanism of IFN-I cDC2 conditioning for T_H_2 responses remains largely unexplored.

In the present study, we used an intradermal immunization model to investigate the role of IFN-I in T_H_2 and TFH responses to allergens. Using DC-specific IFNAR1 conditional KO mice (IFNAR1^ΔCD11c^) and phenotypic characterization of the CD4+ T cell response to allergens, we show that IFN-I signaling in cDC2s increased the differentiation of TFH and T_H_2 cells by increasing CD4+ T cell proliferation following priming. IFNAR1 signaling in cDC2s was also shown to increase the number of *Nb*+ cDC2s in the skin-draining LN without affecting their expression of costimulatory markers. Bulk RNAseq of *Nb*+ CD11b-hi and CD11b-low cDC2s from IFNAR1^WT^ and IFNAR1^ΔCD11c^ mice showed that IFN-I signaling increased the expression of several transcripts including the proliferative and pro-survival cytokine *Il15* (30) and its receptor *Il15ra*. Together, our data demonstrate the importance of IFN-I signaling in DCs for the development of robust CD4+ T cell proliferation and T_H_2 responses to allergens, showing how cDC2s may be programmed by a skin cytokine to prime T_H_2 cells in the dLN.

## Methods

### Mice

All mice were bred and housed at the Malaghan Institute of Medical Research animal facility in specific pathogen-free conditions and were age- and sex-matched within experiments. C57BL/6J and B6-SJL-Ptprca were originally obtained from the Jackson Laboratory (Bar Harbor, ME). *Ifnar1* fl/fl.Itgax-creGFP (IFNAR1^ΔCD11c^) (31, 32) were from Dr. Ashraful Haque, QIMR Berghofer Medical Research Institute, Brisbane, Australia; Cre-negative cage-mates were used as negative controls. 4C13R dual reporter mice (33) were provided by the late Dr. William E. Paul, National Institutes of Health, Bethesda, MD and bred to B6-SJL-Ptprca mice for experiments. All experiments were approved by the Victoria University of Wellington Animal Ethics Committee and performed according to institutional guidelines.

### Immunizations and *in vivo* treatments

*N. brasiliensis* (*Nb*) infective L3 larvae were prepared, washed in sterile PBS and killed by three freeze-thaw cycles as described previously (34, 35). For immunizations, mice were anesthetized and injected intradermally (i.d.) with 300 non-viable *Nb*, 200µg D. pteronyssinus whole bodies (HDM; Greer Laboratories), 4x10^6^ heat-killed *M. smegmatis* (*Ms*; mc2155) or 1x10^7^ heat-killed *C. albicans* (*Ca*; ATCC10231) into the ear pinna. For assessment of responses to 1-Fluoro-2,4-dinitrobenzene (DNFB; Sigma Aldrich), mice were anesthetized and 0.3% DNFB in 1:4 olive oil/acetone was applied epicutaneously to the ear skin. To block IFN-I signaling *in vivo,* mice were treated with 200µg αIFNαR1 (MAR1-5A3) or mouse IgG1 isotype control (MOPC-21) mAb (BioXCell) i.d. at the time of immunization.

### Culture of bone marrow dendritic cells

Mice were euthanized and femurs and tibia were removed. Bone marrows were flushed using a 21G needle and syringe and made into single cell suspensions by pipetting with a 10mL serological pipette before passing through a 70μM sterile filter. Cell suspensions were centrifuged and resuspended in IMDM (Gibco) supplemented with 5% Fetal Calf Serum (Gibco), 1% Penicillin/Streptomycin (Gibco) and 4% Flt3L supernatant from the cell line CHO flag Flk2.clone5 (kindly provided by N. Nicola, WEHI). Cells were plated at 5x10^6^/mL into 6-well plates (ThermoFisher) and cultured at 37°C with 5% CO2 incubator and fed with fresh Flt3L-containing medium every 3 days until maturity at 10 days. BMDCs were processed for confocal microscopy using a 10X expansion microscopy protocol previously described in (36), and imaged using an FV3000 scanning laser confocal microscope (Olympus).

### Adoptive transfer experiments

CD4+ T cells were purified from spleens and LNs of B6-SJL-ptprca donor mice using the Dynabead® FlowComp™ Mouse CD4+ Enrichment kit (Invitrogen) and labelled with CellTrace Violet (ThermoFisher) in sterile PBS for 30 minutes at 37°C. 3 – 5 x 10^6^ labelled cells were then injected intravenously into Ifnar1 fl/fl.Itgax-IFNAR1creGFP recipients. To quantify cell division in adoptively transferred CD4+ T-cells *in vivo*, the following equations were applied to manually gated cell division peaks to generate cell division statistics, where i = division number (undivided cells = 0) and N_i_ = number of events within the division generation (37). These calculations were performed using FlowJo v10 (BD).

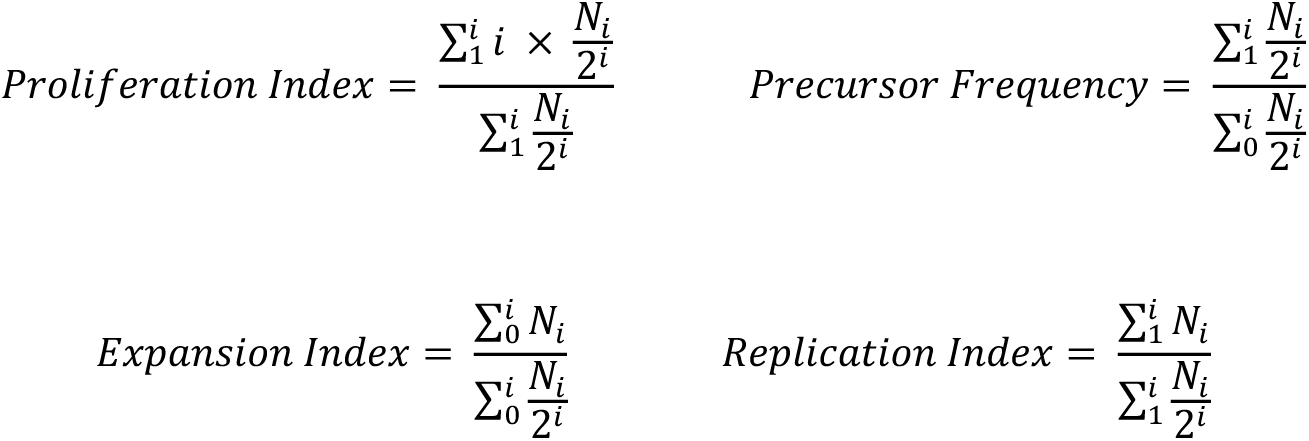

### Cell Preparations

For T cell preparations, LN single cell suspensions were prepared by gently pressing through a 70µm cell strainer using the plunger of a 1mL syringe and rinsing with I-IMDM. For assessment of T cell intracellular cytokines, LN single cell suspensions were cultured in fetal calf serum-supplemented IMDM (Invitrogen) in the presence of 50ng/mL PMA (Sigma-Aldrich), 1µg/mL ionomycin (Merck Millipore) and 1µL/mL GolgiStop™ (BD Pharmingen) for 5 hours at 37°C. cDC were prepared from auricular LNs by digesting with 100µg/mL Liberase TL and 100µg/mL DNase I (both Roche, Germany) for 20 minutes at 37°C before passing through a 70µm cell strainer. Spleens were processed by gently pressing through a cell strainer, and RBCs were lysed using sterile RBC lysis buffer (Sigma) prior to further processing.

### Flow Cytometry

Non-viable cells were stained with Zombie NIR (Biolegend) in PBS. For staining of cell surface markers, cells were suspended in anti-mouse Fc receptors block (clone 2.4G2, affinity purified from hybridoma culture supernatant) prior to labelling with cocktails of fluorescent antibodies made up in PBS containing 2mM EDTA, 0.01% sodium azide and 2% FCS, supplemented with Brilliant Staining Buffer (Biolegend). For staining of TFH cells, cells were incubated with anti-CXCR5 (clone 2G8; BD Pharmingen) for 30 minutes at 37°C prior to staining for additional cell surface antigens. For staining of intracellular or intranuclear antigens, cells were surface stained and then fixed and permeabilized using a FOXP3 Transcription Factor staining kit (ThermoFisher) prior to staining with intracellular antibodies. Antibodies and streptavidin conjugates used in experiments are detailed in Table 1. Compensation or spectral reference controls were recorded using single cell suspensions from relevant tissues, or using UltraComp eBeads™ (ThermoFisher) and dead cells and doublets were excluded from analysis. Samples were collected on a LSRFortessa SORP™ or LSRII SORP™ flow cytometer (both from Becton Dickinson, San Jose, CA), or a 3- or 5-laser Aurora Spectral analyser (Cytek, Fremont, CA) and analysed using FlowJo software (version 10, Becton Dickinson, San Jose, CA).

**Table 1:** Antibodies and Streptavidin Conjugates used in this work.

| Specificity | Clone | Fluorochrome | Company | RRID |
| --- | --- | --- | --- | --- |
| **BCL6** | K112-91 | Alexa Fluor 647 | BD | AB_10898007 |
| **CD3** | 145-2C11 | BUV395 | BD | AB_2738278 |
|  | OKT3 | Alexa Fluor 488 | eBioscience | AB_1907370 |
| **CD4** | GK1.5 | BV750 | Biolegend | AB_2734150 |
|  |  | APC-eFluor 780 | ThermoFisher | AB_11218896 |
|  |  | FITC | BD | AB_395013 |
|  | RM4-5 | BV605 | BD | AB_2687549 |
| **CD8** | 53-6.7 | APC-H7 | BD | AB_1645237 |
|  |  | PerCP Cy5.5 | BD | AB_394081 |
|  |  | Pacific Blue | BD | AB_397029 |
| **CD11b** | M1/70 | BV570 | Biolegend | AB_10896949 |
|  |  | Biotin | BD | AB_394773 |
| **CD11c** | N418 | BV650 | Biolegend | AB_2562414 |
|  | HL3 | Biotin | BD | AB_395059 |
| **CD19** | 6D5 | Alexa Fluor 488 | Biolegend | AB_493339 |
| **CD25** | PC61 | BV480 | BD | AB_2739593 |
|  |  | APC | BD | AB_398623 |
| **CD26 (DPP4)** | H194-122 | PE | Biolegend | AB_2093729 |
| **CD38** | T10 | BV421 | Biolegend | AB_2870503 |
| **CD44** | IM7 | Alexa Fluor 700 | eBioScience | AB_494011 |
|  |  | BUV737 | BD | AB_2870126 |
| **CD45** | 30-F11 | BV510 | Biolegend | AB_2563061 |
| **CD45.1** | A20 | BV510 | Biolegend | AB_2563378 |
|  |  | PE | eBioscience | AB_465675 |
| **CD45R (B220)** | RA3-6B2 | Pacific Blue | BD | AB_397031 |
|  |  | PE-Cy5 | Biolegend | AB_312995 |
|  |  | BUV395 | BD | AB_2738427 |
|  |  | Biotin | eBioScience | AB_466449 |
| **CD74 (CLIP)** | In1 | Alexa Fluor 647 | Biolegend | AB_2632609 |
| **CD80** | 16-10A1 | Biotin | BD | AB_395037 |
| **CD86** | GL1 | BUV395 | BD | AB_2738664 |
| **CD172a (SIRPa)** | P84 | PE-Cy7 | Biolegend | AB_2563546 |
| **CD185 (CXCR5)** | 2G8 | Biotin | BD | AB_394301 |
| **CD279 (PD-1)** | 29F.1A12 | BV785 | Biolegend | AB_2563680 |
| **CD326 (EpcAM)** | G8.8 | PerCP Cy5.5 | Biolegend | AB_2098647 |
| **FcγR block** | 24G2 | Unconjugated | Produced in-house |  |
| **FcER1** | MAR-1 | Biotin | Biolegend | AB_1626100 |
| **FOXP3** | FJK-16S | Alexa Fluor 532 | ThermoFisher | AB_11219265 |
| **GATA3** | L50-823 | Alexa Fluor 647 | BD | AB_1645316 |
| **GL7** | GL7 | FITC | Biolegend | AB_2561697 |
| **IFNγ** | XMG1.2 | BV480 | ThermoFisher | AB_2739549 |
|  |  | APC | BD | AB_398551 |
| IgD | 11-26c.2a | APC-H7 | BD | AB_2739201 |
| IL-4 | 11B11 | APC | BD | AB_395391 |
|  |  | PE | BD | AB_398556 |
| IL-17A | eBio17B7 | PE-Cy7 | ThermoFisher | AB_10732356 |
| Ly6A/E | D7 | PE-CF594 | BD | AB_2737751 |
| Ly6c | HK1.4 | PE-Cy7 | Biolegend | AB_1732093 |
|  |  | Biotin | Biolegend | AB_1236553 |
| Ly6G | 1A8 | Biotin | Biolegend | AB_1186105 |
| MHC-II (IA/IE) | M5/114 | BV605 | BD | AB_2738190 |
|  |  | APC-Cy7 | Biolegend | AB_2069377 |
|  |  | Purified | BD | AB_396545 |
|  | 3JP | Alexa Fluor 488 | Produced in-house |  |
|  |  | Alexa Fluor 647 | Produced in-house |  |
|  |  | Purified | Produced in-house |  |
| Streptavidin | N/A | BUV737 | BD | AB_2869560 |
|  |  | PE-CF594 | BD | AB_11154598 |
|  |  | PE-Texas Red | BD | AB_10054235 |
| TcRβ | H57-597 | BUV395 | BD | AB_2740818 |
|  |  | BV605 | BD | AB_2687544 |
| TcRgd | GL3 | BV605 | Biolegend | AB_2563356 |
| XCR1 | ZET | BV421 | Biolegend | AB_2565230 |

### Staining of surface and intracellular MHCII

For the analysis of surface and intracellular MHC-II molecule expression, following extracellular MHC-II staining, remaining unbound surface MHC-II molecules were saturated using unconjugated purified anti-MHC-II M5/114 (BC) or 3JP (made in-house). Cells were then fixed, permeabilized, and stained intracellularly for MHC-II using the same antibody clone in different fluorochromes.

### Cell sorting

Sterile single cell suspensions were prepared as detailed above and incubated in FACS tubes with Zombie NIR in PBS for 25 minutes on ice, followed by 2.4G2 Fc block for 10 minutes on ice in PBS supplemented with 5% FCS. Cells were spun down and incubated with sterile sorting antibody master mix for 15 minutes on ice.

Cell sorting was performed using a BD Influx® cell sorter or Cytek Aurora CS. Machine cleaning, laser alignment and QC were performed on the day of the sort, with manual compensation or spectral unmixing performed using single stained cells or beads.

### Quantitative RT-PCR

RNA was isolated from cell lysates using a Zymo quick-RNA microprep kit (Zymo Research). RNA samples were then immediately converted to cDNA using a High Capacity RNA-to-DNA kit (Applied Biosystems), and samples were then stored at -20°C before further use. cDNA was preamplified using relevant TaqMan probes (ThermoFisher) in Sso Advanced Preamplifiation mix (Bio-Rad) for 12 cycles, and qPCR reactions performed in 384-well plates using TaqMan Universal Master Mix (applied biosystems), in a QuantStudio Flex 7 RealTime qPCR Machine for 40 cycles (Applied Biosystems). TaqMan probes used in this study are detailed in Table 2.

**Table 2:**
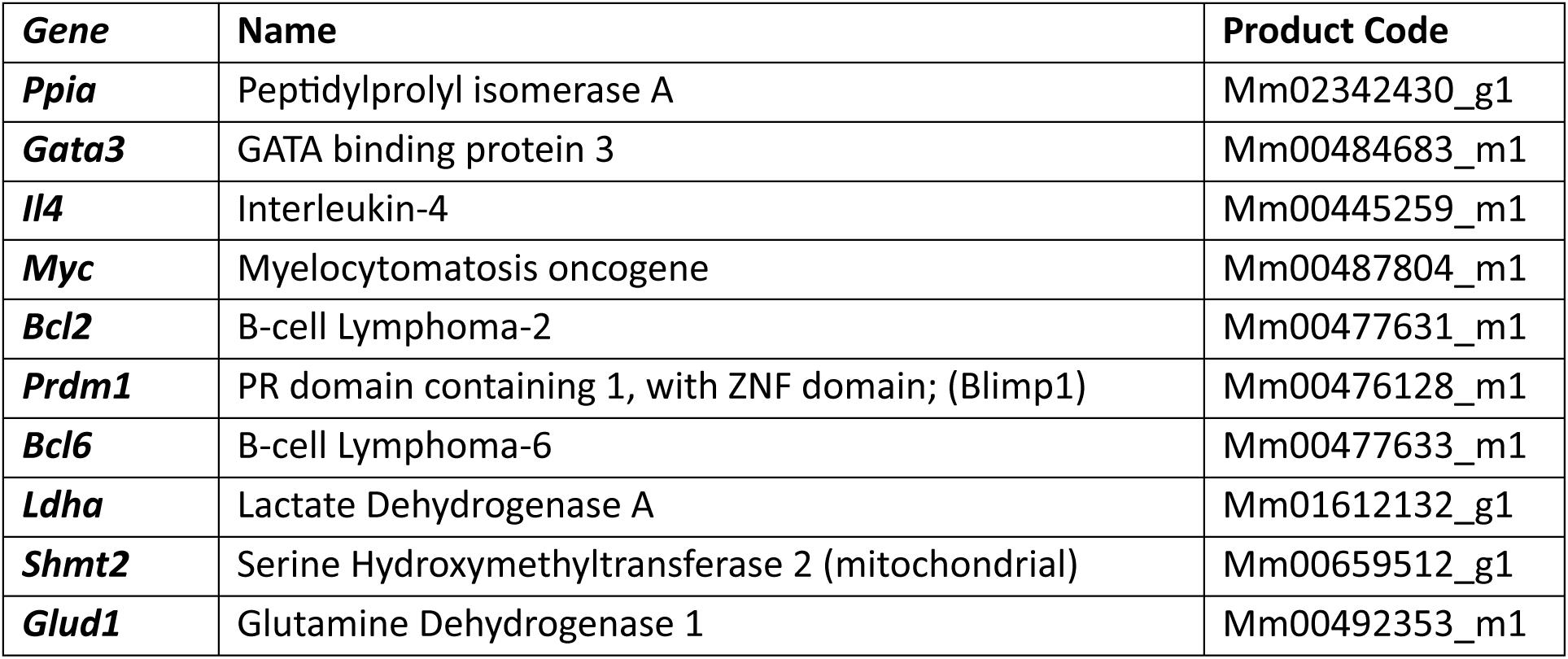
TaqMan qPCR probes used in this work.

### RNA isolation and sequencing

Cells were sorted directly into QIAzol Lysis buffer, and frozen at -70°C. RNA was extracted from cell lysates using a miRNeasy kit (QIAgen), and resuspended in nuclease-free water. RNA sequencing was contracted to Otago Genomics NZ. Sample QC was performed using a Fragment Analyzer Systems (Agilent), and libraries prepared using a SMARTer ® Stranded Total RNA-Seq Kit v3 – Pico Input Mammalian (Takara). Samples were then sequenced using an Illumina NextSeq 2000 (Illumina) using Illumina TruSeq kits.

### Read mapping and DEG analysis

Bulk RNA-sequencing data was provided by Otago Genomics (NextSeq 2000), and Illumina basespace. Reads were trimmed using cutadapt version 4.9 (38), and aligned to mouse genome M35 GRCm39 (39) using STAR version 2.7.11b (40). Aligned reads were counted using R package Rsubread featureCounts (41). Differentially expressed genes were computed using R package DESeq2 (42), and are listed in supplementary files. Data exploration, graphing, and analysis was performed using in-house R Shiny app (43). Raw data, source code, and singularity recipe files are available at **10.5281/zenodo.13698207**.

### ELISA

Mouse serum IgE was quantified using a ThermoFisher uncoated ELISA kit in 96-well Nunc MaxiSORP plates. *Nb*-specific IgG1 titres were measured using an in-house developed assay. 96-well Nunc MaxiSORP plates were coated overnight with *Nb* homogenates at 5ng/mL. The following day, plates were washed with PBS-Tween and blocked with 100uL of PBS-BSA at 5%, before being washed with PBS-Tween. Mouse serum dilutions were then added to relevant plates and incubated for an hour, before being washed again with PBS-Tween and incubated with serum samples following washing with PBS + 0.05% Tween-20. Plates were incubated with 100uL of anti-IgG1 Horseradish Peroxidase (HRP) conjugated antibody for an hour and washed again. Plates were then incubated with 100uL TMB substrate solution (BD) and left to develop for 15 minutes before 100uL of 2N H_2_SO_4_ was added to stop the reaction. *Nb*-specific IgG1 antibody affinity index was determined by Urea-dissociation assay, duplicate samples were incubated with 6M Urea at 37°C. All ELISA plates were read using a TECAN plate reader, measuring absorbance at 450nm.

### Statistical analyses

Statistical analyses were performed using Prism 10 GraphPad software. Comparisons between 3 or more groups were by one-way ANOVA with Holm-Šidak’s correction for multiple comparisons. When more than one variable was assessed, two-way ANOVA with Šidak’s correction was used. Comparisons of two groups only were made with Student’s t-test, or when sample sizes were small or not normally distributed, using Mann-Whitney tests. P values lower than 0.05 were considered significant and are referred to as follows: **** = p < 0.0001, *** = 0.0001 < p < 0.001, ** = 0.001 < p < 0.01, * = 0.01 < p < 0.05, ns = not significant.

## Results

### IFN-I signaling in DCs regulates CD4+ T cell responses to allergens

Previous studies have shown a role for IFN-I signaling in DCs in the regulation of T_H_2 differentiation after immunization with *Nb*, HDM and SEA (12, 13, 44) but lacked a characterization of the T_H_2 response. We investigated cytokine responses to two allergens, the inactivated L3 larvae of the rodent hookworm *Nb,* or whole bodies of the house dust mite *Dermatophagoides pteronyssinus*. Mice were injected intradermally (i.d.) with *Nb* or HDM and IFNAR1 signaling was blocked at the time of immunization by i.d. treatment with aIFNAR1 antibodies or IgG1 isotype control. Immunized mice treated with aIFNAR1 displayed reduced numbers of IL-4+ (**Fig 1A** and **S1A**) and IFNγ+ CD4+ T cells (**Fig 1B**) when compared to isotype-treated mice. The number of CD44-hi CD4+ T cells was also significantly reduced by aIFNAR1 treatment (**Fig 1C**), indicating that IFNAR1 signaling was required for the upregulation of T cell activation markers as well as cytokine expression. By contrast, mice immunized with heat-killed *Mycobacterium smegmatis* (*Ms*) or treated epicutaneously with DNFB generated similar numbers of IFNγ+ and CD44-hi CD4+ T cells regardless of aIFNAR1 treatment (**Fig S1B – E).** Similarly, treatment with aIFNAR1 did not affect the numbers of CD44-hi and IL-17A+ CD4+ T cells in mice immunized with heat-killed *Candida albicans* (*Ca*) (**Fig S1F, G**), indicating that IFN-I signaling was not required for the development of responses to bacterial or yeast antigens in this context.

**Figure 1:**
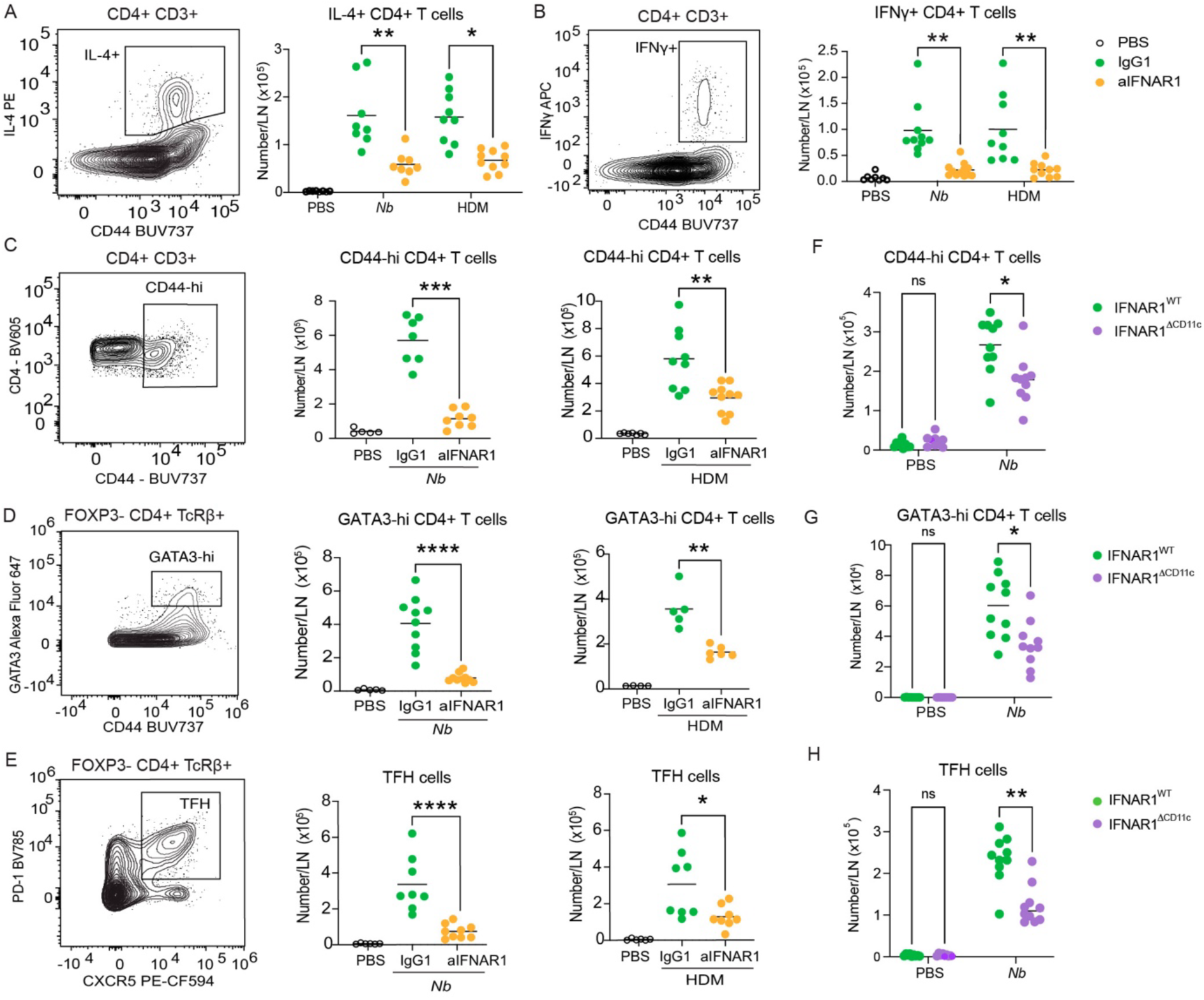
IFN-I signalling in CD11c+ cells is necessary for T_H_2 and TFH differentiation in LN after allergen immunization. **A-E:** C57BL/6 mice were immunized intradermally with HDM or *Nb* and treated with aIFNAR1 or isotype control. T cell responses in the skin-draining LN were assessed by flow cytometry 7 days following immunization. **A:** Representative gating and numbers of IL-4+ CD4+ T cells. **B:** Representative gating and numbers of IFNγ CD4+ T cells. **C:** Representative gating and numbers of CD44-hi CD4+ T cells. **D:** Representative gating and numbers of GATA3-hi FOXP3-CD4+ T cells. **E:** Representative gating and numbers of FOXP3-TFH. **F-H:** IFNAR1^WT^ and IFNAR1^ΔCD11c^ mice were immunized intradermally with HDM or *Nb*, and the skin-draining LN response was assessed by flow cytometry 5 days following immunization. **F:** Numbers of CD44-hi FOXP3-CD4+ T cells. **G:** Numbers of GATA3-hi FOXP3-cells. **H:** Numbers of FOXP3-TFH. Dot plots show data from two pooled experiments each with 3 – 5 mice per group. Each dot corresponds to one mouse. *P* values refer to comparisons between the indicated groups and were calculated using Kruskall-Wallis test in **A, B**; Mann-Whitney tests in **C, E**, **G**; and two-way ANOVA in **D, F**, **H**. ****, *p*<0.0001; ***, *p*<0.001; **, *p*<0.01; *, *p*< 0.05; ns, *p*≥0.05.

As IL-4 is made in the dLN by both T_H_2 and TFH cells (45, 46), we asked whether both populations required IFNAR1 signaling. The T_H_2 and TFH populations were identified on the basis of high GATA3 and CXCR5 expression, respectively, and showed minimal overlap by flow cytometry (**Fig S2A**). Following immunization with *Nb* or HDM, there were fewer GATA3-hi T_H_2 cells in aIFNAR1-treated mice compared to those treated with isotype control (**Fig 1D**). In addition, mice treated with aIFNAR1 generated fewer TFH in response to *Nb* and HDM compared to isotype controls (**Fig 1E**).

Treatment with aIFNAR1 blocks IFNAR1 signaling in all cells. To restrict the involvement of IFN-I signaling to CD11c+ cells (which include cDCs, pDCs, Langerhans cells and some monocytes), we assessed the T_H_2 response to *Nb* in mice with a Cre-mediated deletion of IFNAR1 in CD11c+ cells (IFNAR1^ΔCD11c^). Similar to aIFNAR1-treated mice, IFNAR1^ΔCD11c^ mice generated lower numbers of CD44-hi, GATA3-hi and TFH CD4+ T cells compared to IFNAR1^WT^ mice (**Fig 1F – H**), indicating that IFN-I signaling in CD11c+ cells is necessary for optimal CD4+ T cell responses to these T_H_2 antigens. The numbers of total FOXP3+ regulatory T (Treg) cells were similar in both strains (**Fig S2B**). By contrast, the numbers of GATA3-hi and CXCR5+ Tregs (**Fig S2C, D**) were higher in IFNAR1^WT^ mice compared to IFNAR1^ΔCD11c^.

These results suggest that IFNAR1 signaling in CD11c+ DCs is necessary for optimal Th2 and TFH responses after immunization with Th2 antigens.

### TFH differentiation requires IFN-I signaling in cDCs

To further characterize how IFN-I signaling supports the development of TFH responses to *Nb*, we utilized 4C13R *Il4*-AmCyan (*Il4*-AmC), *Il13*-DsRed dual reporter mice to enable the direct assessment of *Il4*-AmC+ cells *ex vivo* without the need for *in vitro* restimulation (46) (**Fig 2A**). Mice were immunized with *Nb* and treated with aIFNAR1 or isotype, and the draining LN response was assessed 5 days following immunization. Compared to isotype controls, mice treated with aIFNAR1 displayed reduced numbers of *Il4*-AmC+ CD4+ T cells (**Fig2B**). The number of *Il4*-AmC+ TFH was also significantly reduced in mice given IFNAR1 blockade (**Fig 2C**) without affecting the median FI of *Il4*-AmC, suggesting that the function of the remaining TFH was likely preserved.

**Figure 2:**
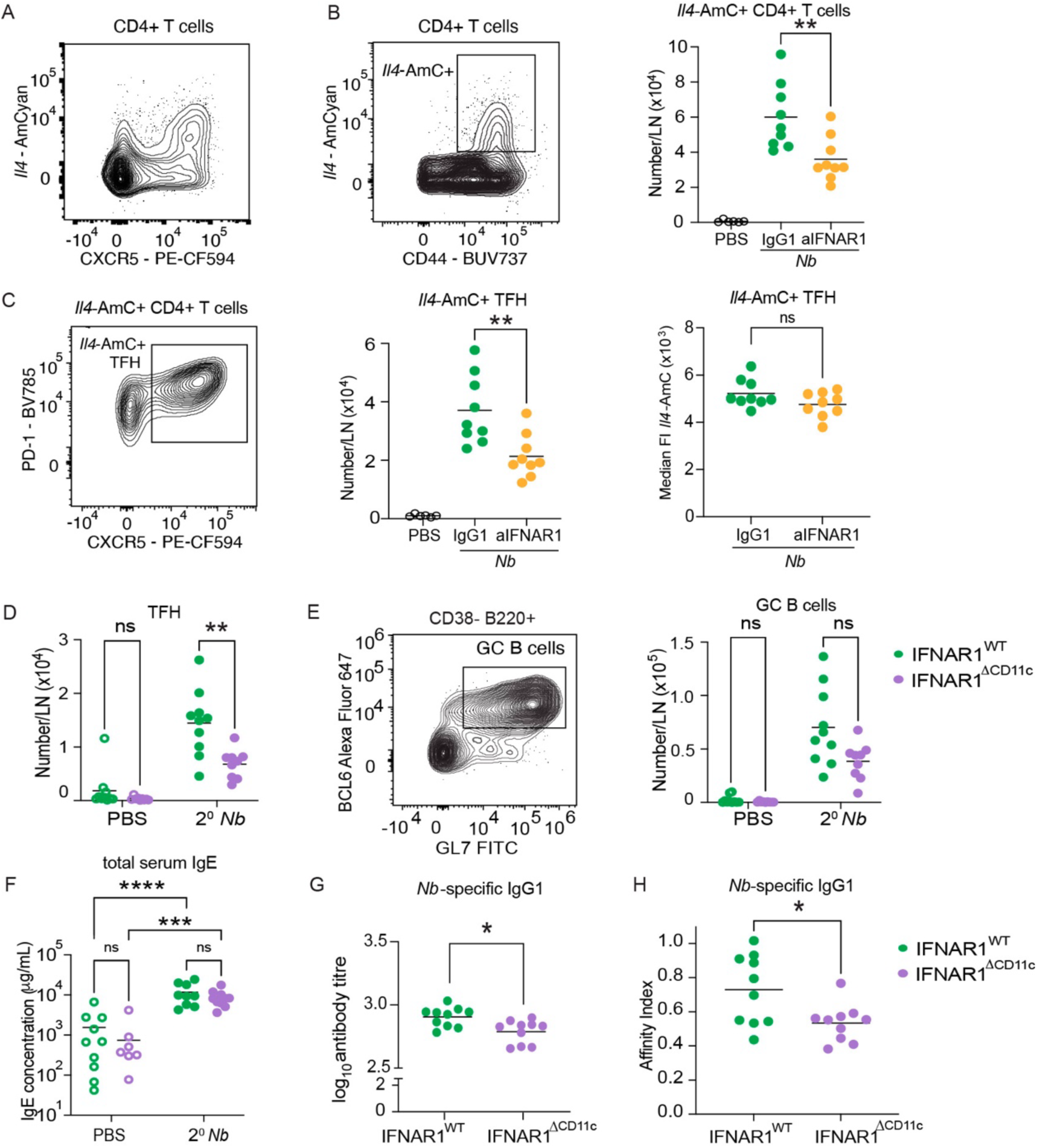
IFN-I signalling is essential for TFH and IgG1 responses to *Nb*. **A-C:** 4C13R mice were immunized intradermally with *Nb* and treated with aIFNAR1 or isotype control on day 0. T cell responses in the skin-draining LN were assessed by flow cytometry on day 5. **A:** Representative contour plot showing of *Il4*-AmC and CXCR5 expression in CD4+ T cells. **B:** Representative gating and numbers of *Il4*-AmC+ CD4+ T cells. **C:** Representative gating and numbers of *Il4*-AmC+ TFH, and *Il4-*AmC Median fluorescence intensity (MFI) in *Il4*-AmC+ TFH. **D-H**: IFNAR1^WT^ and IFNAR1^ΔCD11c^ mice were immunized intradermally with *Nb* on day 0 and day 21. The T and B cell responses in draining LN and antibody responses in serum were assessed on day 28. **D:** Numbers of TFH. **E:** Representative gating and number of GC B cells. **F:** Total serum IgE. **G:** *Nb*-specific IgG1 titre. **H:** *Nb*-specific IgG1 affinity. Dot plots show data from two pooled experiments each with 4 – 5 mice per group. Each dot corresponds to one mouse. *P* values refer to comparisons between the indicated groups and were calculated using Mann-Whitney tests in **B, C, G**, **H;** Multiple Mann-Whitney tests with Holm-Šidak’s correction in **D, E;** and two-way ANOVA in **F**. ****, *p*<0.0001; ***, *p*<0.001; **, *p*<0.01; *, *p*< 0.05; ns, *p*≥0.05.

To further assess the TFH response in the absence of IFN-I signaling in DCs, IFNAR1^ΔCD11c^ mice were immunized with *Nb* and given a second *Nb* injection 3 weeks later. CD4+ T cell and B cell responses in the skin-draining LN and spleen were assessed one week following the second treatment. Twice-immunized IFNAR1^ΔCD11c^ mice displayed a reduced frequency of LN TFH compared to IFNAR1^WT^ (**Fig 2D**). The frequency of TFH in spleen did not increase after immunization and was similar in IFNAR1^WT^ and IFNAR1^ΔCD11c^ mice (**Fig S**). Despite reduced TFH cell numbers in IFNAR1^ΔCD11c^ compared to IFNAR1^WT^ mice, there was no concurrent defect in the frequencies of LN germinal centre B cells (**Fig 2E**), and no decrease in total serum IgE (**Fig 2F**). However, serum *Nb*-specific IgG1 titres and affinity index were significantly reduced in IFNAR1^ΔCD11c^ mice compared to IFNAR1^WT^ (**Figure 2G, H**), likely due to the lower frequency of TFH leading to a reduction in antibody affinity maturation. Taken together, these results show that IFNAR1 signaling in DCs is required for TFH differentiation after *Nb* immunization.

### IFN-I signaling in cDCs facilitates the expansion of CD4+ T-cells during T_H_2 responses

To further assess the impact of IFN-I signaling in DCs on TFH and T_H_2 differentiation, we developed an adoptive transfer model to assess CD4+ T cell proliferation *in vivo*. CD45.1+ polyclonal CD4+ T cells were labelled with CellTrace Violet (CTV) and injected i.v. into IFNAR1^ΔCD11c^ and IFNAR1^WT^ recipients. Recipient mice were immunized with *Nb* 24h later, and cell division in the skin-dLN was measured on day 5 after immunization (**Fig 3A**). The frequency of total CD4+ donor cells was not significantly different between IFNAR1^ΔCD11c^ and IFNAR1^WT^ mice (**Fig 3B**). However, IFNAR1^ΔCD11c^ mice displayed significantly reduced numbers of CD4+ T cells at division peak 10+ compared to IFNAR^WT^ mice, suggesting a reduced proliferative and/or survival capacity (**Fig 3C**). The frequency of GATA3-hi and TFH cells within the division 10+ subset were not significantly different between IFNAR1^WT^ and IFNAR1^ΔCD11c^, indicating that both subsets experienced a similarly defective expansion in IFNAR1^ΔCD11c^ mice (**Fig 3D**). Although there was no significant difference between IFNAR1^WT^ and IFNAR1^ΔCD11c^ in average number of divisions undergone by responding cells after immunization (**Fig 3E**), the fold-expansion of dividing CD4+ T cells was higher in IFNAR1^WT^ compared to IFNAR1^ΔCD11c^ mice (**Fig 3E**).

**Figure 3:**
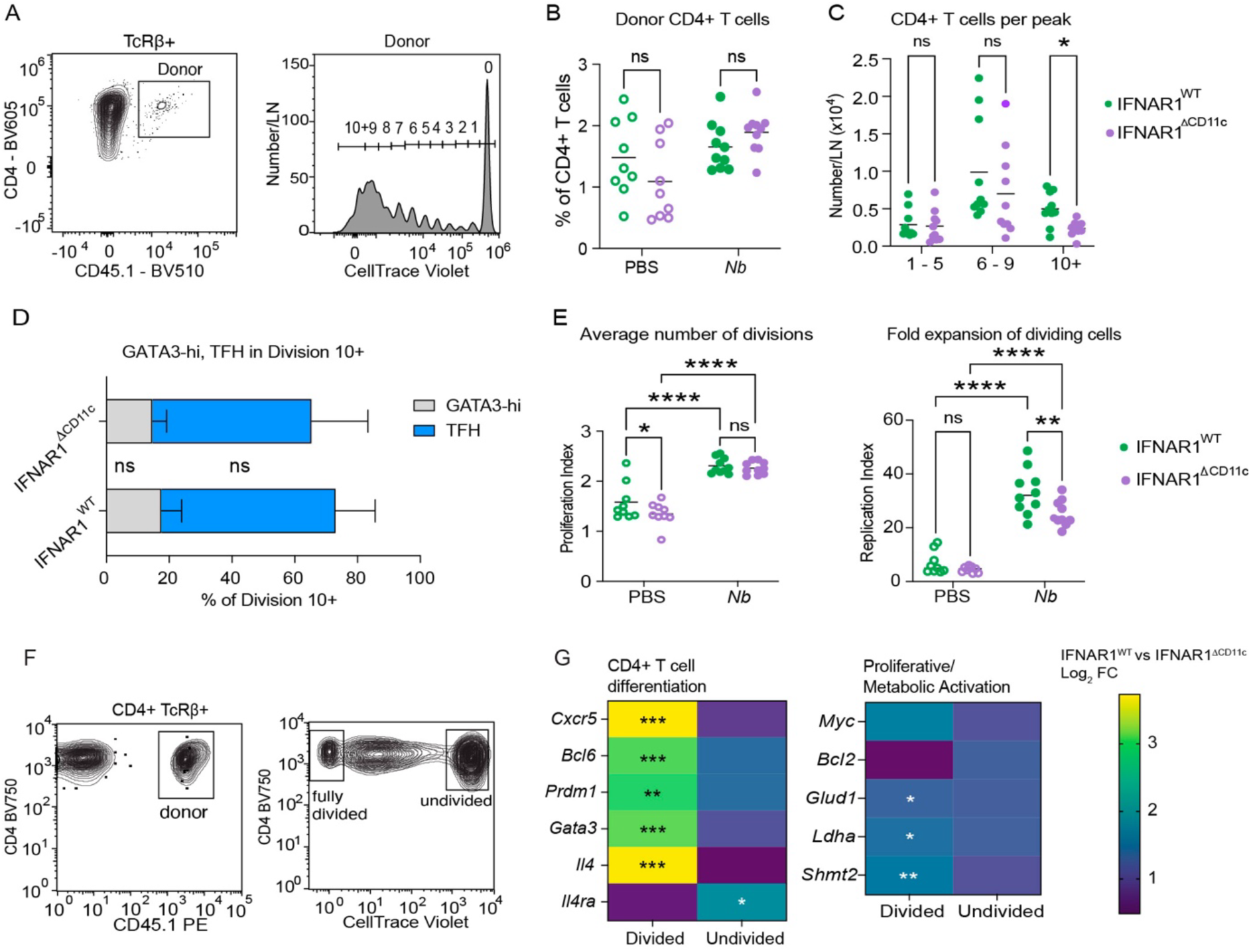
IFN-I signalling in CD11c+ cells facilitates CD4+ T cell expansion in response to allergens. IFNAR1^WT^ and IFNAR1^ΔCD11c^ mice were adoptively transferred with 3 x10^6^ CTV-labelled polyclonal CD4+ T cells, and immunized intradermally with *Nb* 24 hours following adoptive transfer. The CD4+ T cell response in draining LN was assessed by flow cytometry (**A -E**) or RT-qPCR (**F, G**) 5 days following immunization. **A:** Representative gating of donor CD4+ T cells and division peaks as measured by CellTrace Violet dilution **B:** Frequencies of donor CD4+ T cells. **C**: Numbers of donor CD4+ T cells in the indicated CTV division peaks. **D:** Frequencies of GATA3-hi cells and TFH in the 10+ division peak. **E:** Average Number of Divisions (Proliferation Index) and Fold expansion (Replication Index) of divided CD4+ T cells as estimated by CTV dilution. IFNAR1^WT^ and IFNAR1^ΔCD11c^ mice were given 3 x10^6^ adoptively transferred CTV-labelled polyclonal CD4+ T cells i.v., and immunized with *Nb* 24 hours following adoptive transfer. **F:** Representative gating for sorting of undivided and fully divided donor CD4+ T cells in *Nb*-immunized mice. **G:** Expression of selected genes as determined by RT-qPCR. Data are expressed as Log2 fold-change of the mean normalized expression of each gene. Dot plots show data from two pooled experiments each with 5 mice per group. Each dot corresponds to one mouse. *P* values refer to comparisons between the indicated groups and were calculated using multiple Mann-Whitney tests with Holm-Šidak’s correction in **B - E**, and multiple Mann Whitney tests without correction in **F**. ****, *p*<0.0001; ***, *p*<0.001; **, *p*<0.01; *, *p*< 0.05; ns, *p*≥0.05.

To further investigate the characteristics of dividing CD4+ T cells in IFNAR1^ΔCD11c^ and IFNAR1^WT^ mice, we compared gene expression in FACS-sorted undivided and fully divided donor CD4+ T cells (**Fig 3F**) using RT-qPCR. Compared to IFNAR1^WT^, fully divided cells from IFNAR1^ΔCD11c^ mice displayed a significantly lower expression of key TFH and T_H_2 effector differentiation genes including *Cxcr5*, *Bcl6*, *Prdm1*, *Gata3*, and *Il4* (**Fig 3G**). In addition, *Il4ra* expression was lower in undivided cells from IFNAR1^ΔCD11c^ mice compared to IFNAR1^ΔCD11c^, presumably due to lower IL-4 production and signaling in IFNAR1^ΔCD11c^ LN (**Fig 3G**) (47, 48, 49). While there was no significant difference in the expression of the cell division timer *Myc* or apoptotic regulator *Bcl2* (50) between divided CD4+ T cells from IFNAR1^ΔCD11c^ and IFNAR1^WT^ mice, IFNAR1^ΔCD11c^ mice displayed lower expression of genes involved in amino acid (*Glud1, Shmt2*) and lactic acid (*Ldha*) metabolism (**Fig 3F**), suggesting lower metabolic activity in divided cells from IFNAR1^ΔCD11c^ mice.

### IFN-I signaling in migratory cDC2s is not required for the upregulation of costimulatory molecules

To establish how IFNAR1 signaling in cDCs supported Th2 proliferation, we characterized cDC2s in the skin-draining LN following immunization with *Nb.* IFNAR1^WT^ and IFNAR1^ΔCD11c^ mice were immunized with CellTracker Orange-labelled *Nb* (*Nb*-CTO), and the number of *Nb*-CTO+ cDC2s in the skin-draining LN was determined by flow cytometry (**Fig 4A**, gated as in **S4A**). The number of *Nb*-CTO+ CD11b-hi and -low cDC2s was significantly higher in IFNAR1^WT^ compared to IFNAR1^ΔCD11c^ (**Fig 4B**). The expression of the IFN-I response markers BST2 (Tetherin) and Ly6A/E (Sca1) was higher in cDC2s from *Nb*-CTO-immunized IFNAR1^WT^ mice compared to IFNAR1^ΔCD11c^ (**Fig 4C, D**), indicating that IFNAR1 signaling was successfully blocked. However, there was no significant difference in the expression of the costimulatory markers CD80 or CD86 between cDC2s from *Nb*-immunized IFNAR1^WT^ and IFNAR1^ΔCD11c^ mice (**Fig 4E, F**), indicating that the differences in CD4+ T cell expansion cannot be explained by lack of costimulation.

**Figure 4:**
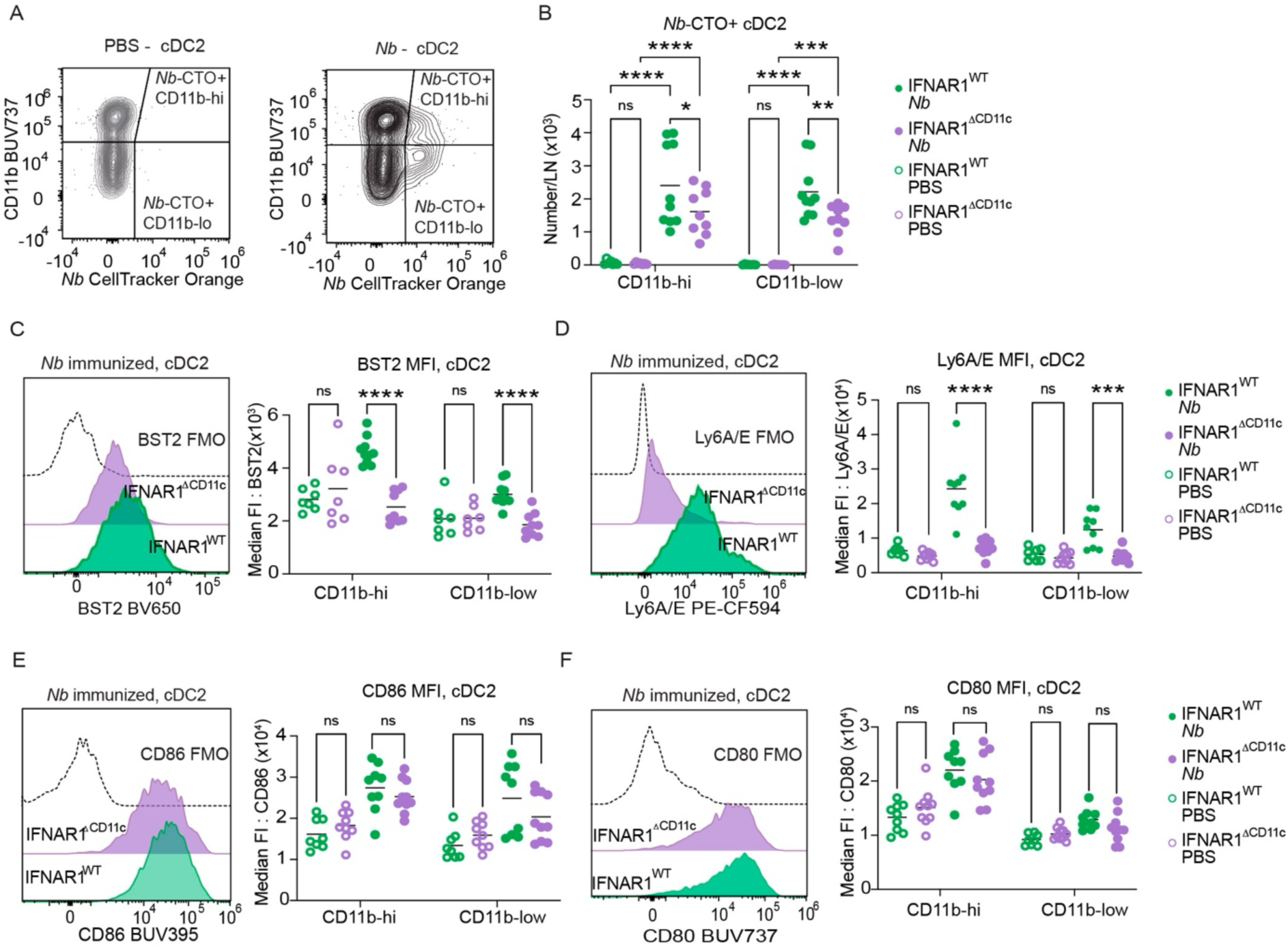
IFN-I signalling in cDC2s regulates CD4+ T cell expansion independently of costimulation. IFNAR1^WT^ and IFNAR1^ΔCD11c^ mice were immunized intradermally with either (**A-B**) CTO-labelled *Nb* (CTO-*Nb*), or (**C-F**) *Nb*, and the skin-draining LN was assessed by flow cytometry two days following immunization. **A:** Representative gating of *Nb*-CTO+ CD11b-hi and CD11b-low cDC2s. **B:** Numbers of *Nb*-CTO+ CD11b-hi and CD11b-low cDC2s. **C:** Representative histograms of BST2 expression in total migratory cDC2s and Median Fluorescence Intensity (MFI) of BST2 expression in total CD11b-hi and CD11b-low cDC2s. **D:** Representative histograms of Ly6A/E expression in total migratory cDC2s and MFI of Ly6A/E expression in total CD11b-hi and CD11b-low cDC2s. **E:** Representative histograms of CD86 expression in total migratory cDC2s and MFI of CD86 expression in total CD11b-hi and CD11b-low cDC2s. **F:** Representative histograms of CD80 expression in total migratory cDC2s and MFI of CD80 expression in total CD11b-hi and CD11b-low cDC2s. Dot plots show data from two pooled experiments each with 4 – 5 mice per group. Each dot corresponds to one mouse. *P* values refer to comparisons between the indicated groups and were calculated using two-way ANOVA. ****, *p*<0.0001; ***, *p*<0.001; **, *p*<0.01; *, *p*< 0.05; ns, *p*≥0.05.

Bone-marrow derived DCs treated with IFN-I were reported to express lower surface MHC-II compared to untreated, increasing their MHC-II turnover following maturation (51). To investigate MHC-II protein turnover, we investigated the surface and intracellular expression of MHC-II on cDC2s from IFNAR1^WT^ or IFNAR1^ΔCD11c^ mice following immunization with *Nb* (**Fig S4C**). No significant differences were observed between cDC2 from IFNAR1^WT^ and IFNAR1^ΔCD11c^ mice in the expression of MHC-II at the cell surface, internalized in intracellular compartments, or newly synthesized and associated with CD74, by flow cytometry (**Fig S4B, Fig S4D - F).** Therefore, in this *in vivo* model, IFN-I signaling does not appear to alter the expression and distribution of MHC-II molecules in cDC2 from *Nb-*immunized or PBS-treated mice.

### Bulk RNAseq analysis of CTO-*Nb*+ cDC2s from IFNAR1^WT^ or IFNAR1^ΔCD11c^ mice reveals differential expression of genes involved in CD4+ T cell proliferation and DC migration

As expression of costimulatory markers and MHC-II compartmentalization were not significantly different between IFNAR1^ΔCD11c^ and IFNAR1^WT^ cDC2s, and to identify potential causes of their different migration and T cell stimulatory ability, we performed a bulk RNAseq transcriptomic characterization of *Nb*+ CD11b-hi and CD11b CD11b-low cDC2s purified from the skin-draining LN 48h after *Nb* injection. Total CD11b-hi and CD11b-low cDC2s from mice treated intradermally with PBS were used as controls (**Fig S5A**). Principal Component Analysis (PCA) of the top 10,000 most variable genes revealed that sorted cDC2s separated according to subset (PC1) and immunization (PC2). However, there was no obvious separation of cDC2s by genotype even at PC3 and PC4, suggesting only subtle differences between IFNAR1^WT^ and IFNAR1^ΔCD11c^ cDC2s (**Fig S5B**). Genes differentially expressed in cDC2s from IFNAR1^ΔCD11c^ compared to IFNAR1^WT^ mice following PBS treatment were very few (**Fig S5C, D**), indicating low to no IFNAR1 signaling in LN cDC2s at steady state.

Differential gene expression analysis of cDC2s from *Nb*-CTO+ cDC2s compared to PBS showed that, in agreement with our flow cytometry data (**Fig 4E, F**), costimulatory marker transcripts were similarly upregulated in both IFNAR1^ΔCD11c^ and IFNAR1^WT^ cDC2 subsets following *Nb* immunization (**Fig S5E, S5F**). Expression of cytokine transcripts was also similar between IFNAR1^WT^ and IFNAR1^ΔCD11c^, apart from *Il12b* which was slightly lower in CD11b-low cDC2 from IFNAR1^WT^ compared to IFNAR1^ΔCD11c^ mice (**Fig S5G, H**).

A comparison of all differentially expressed genes (DEGs) in IFNAR1^WT^ vs IFNAR1^ΔCD11c^ cDC2s following *Nb* immunization revealed 149 transcripts that were significantly higher in IFNAR1^WT^ CD11b-hi cDC2s, and 99 that were higher in IFNAR1^ΔCD11c^ (**Fig 5A**). In CD11b-low cDC2s, 111 transcripts were higher in IFNAR1^WT^ and 31 were higher in IFNAR1^ΔCD11c^ (**Fig 5A**). Only 46 DEGs were in common between both cDC2 subsets (**Fig 5B**), suggesting that the cDC2 response to IFNAR1 signaling following *Nb* immunization is subset specific.

**Figure 5:**
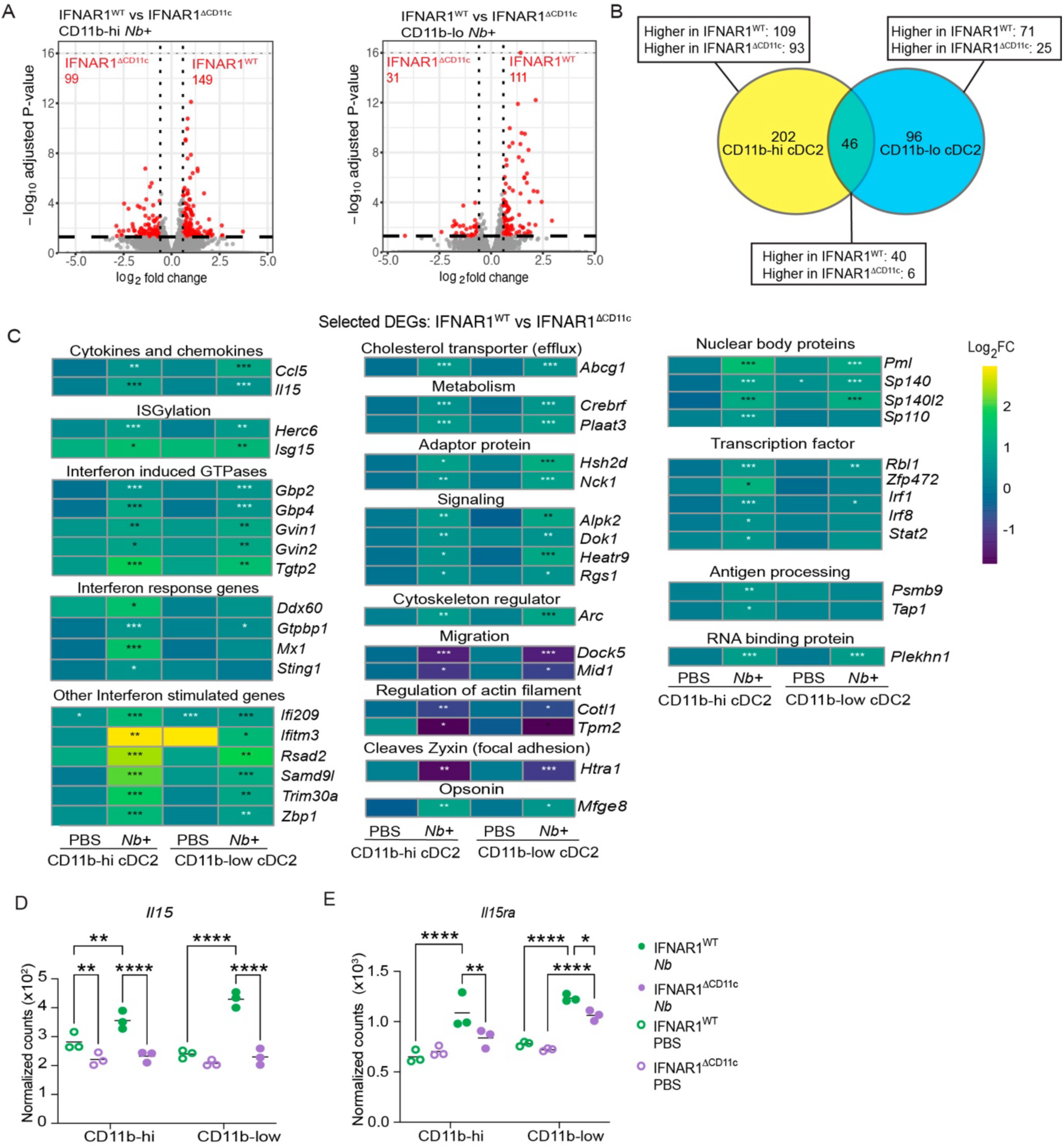
Transcriptomic analysis of CD11b-hi and CD11b-low cDC2s following *Nb* immunization in IFNARWT and IFNARΔCD11c mice. IFNAR1^WT^ and IFNAR1^ΔCD11c^ mice were immunized intradermally with *Nb*-CTO. Draining LN *Nb*-CTO+ CD11b-hi and CD11b-low cDC2s were sorted on day 2 and analysed by bulk RNA sequencing. **A:** Volcano plots showing differential gene expression analysis of IFNAR1^WT^ *vs*. IFNAR1^ΔCD11c^ CD11b-hi and CD11b-low cDC2s. **B:** Venn diagram showing DEG numbers in CD11b-hi and CD11b-low cDC2s in IFNAR1^WT^ *vs* IFNAR1^ΔCD11c^ mice. **C:** Heatmap showing the Log2 fold-change of selected IFNAR1^WT^ vs IFNAR1^ΔCD11c^ DEGs in CD11b-hi and CD11b-low cDC2s. **D:** Normalized feature counts of *Il15* and **E:** *Il15Ra* transcripts in CD11b-hi and CD11b-low cDC2s from IFNAR1^WT^ and IFNAR1^ΔCD11c^ mice. Dot plots show data from three biological replicates each corresponding to 2 – 3 pooled mice per group. Each dot corresponds to one replicate. *P* values in **C** refer to the indicated comparisons and were calculated using the DESeq2 Wald test. ***, *p*<0.001; **, *p*<0.01; *, *p*< 0.05. *P* values in **D – E** refer to the comparisons between the indicated groups and were calculated using two-way ANOVA. ****, *p*<0.0001; ***, *p*<0.001; **, *p*<0.01; *, *p*< 0.05; ns, *p*≥0.05.

We selected DEGs that were upregulated by IFN-I in both cDC2 subsets or were specific to CD11b-low cDC2 (**Fig 5C**) due to the key role of this cDC subset in T_H_2 differentiation (3, 4). These DEGs included only two cytokines, *Ccl5* and *Il15*. CCL5 production by cDCs in LN was shown to recruit monocytes by binding to CCR5 (52). Indeed, monocytes were found to be present in higher numbers in the draining LN of IFNAR1^WT^ compared to IFNAR1^ΔCD11c^ mice following *Nb* immunization (**Fig S5I**). *Il15* has a demonstrated role in T cell proliferative responses and requires trans-presentation to T cells via *Il15ra* (53), which was also more highly expressed on both cDC2 subsets from IFNAR1^WT^ vs IFNAR1^ΔCD11c^ mice (**Fig 5D**). The IFN-regulated *Isg15*, when secreted, can bind LFA-1 and lead to the secretion of IFNγ (54). Other transcripts included interferon-induced GTPases, signal transduction adaptor proteins, intracellular phospholipases, signaling proteins, and nuclear body proteins (**Fig 5C**). The cholesterol transporter *Abcg1* was also upregulated, pointing towards a potential role of IFNAR1 signaling in cDC2 lipid homeostasis (**Fig 5C**). Genes involved in proteasomal antigen processing were also highly expressed in IFNAR1^WT^ compared to IFNAR1^ΔCD11c^ *Nb*-CTO+ CD11b-low cDC2s (**Fig 5C**), which is consistent with their known regulation by IFNAR1 signaling (22).

Consistent with previous literature, the IFN-I-induced transcription factor *Irf1* was lower in IFNAR1^ΔCD11c^ compared to IFNAR1^WT^ in both cDC2 subsets (55), while *Irf8* (17) and *Stat2* (20) were lower only in CD11b-low cDC2s, as was *Sting1* (56) (**Fig 5C**). Four of the genes downregulated in both IFNAR1^WT^ cDC2 subsets were involved in cytoskeletal organization and motility, *Dock5* (57), *Mid1* (58), *Cotl1* (59), and *Tpm2* (60). *Htra1* (61), has a role in the cleavage of Zyxin and could also play a role in cell motility (62). The role of these genes in cDC motility has not been extensively studied, and therefore their role in cDC migration to the LN following *Nb* immunization is largely speculative.

## Discussion

In the present study, we characterize the impact of IFN-I signaling in cDC2s on CD4+ T cell responses to allergens. We show that cDC2s exposed to IFNAR1 signaling primed CD4+ T cells that were able to divide more and expressed higher levels of transcripts characteristic of T_H_2 and TFH differentiation including *Gata3, Bcl6* and *Il4*, thereby resulting in T_H_2 responses of significantly higher magnitude. IFN-I-responsive cDC2s did not express higher costimulatory molecules or cytokines associated with T_H_2 priming, but upregulated *Il15,* a cytokine reported to induce IL-4 production in cultured CD4+ T cells (63{Borger, 1999 #58)}

### IFNAR1 signaling in cDC2s programmes greater CD4+ T cell responses to allergens

Experiments using IFNAR1 blockade and IFNAR1^ΔCD11c^ mice demonstrated that loss of IFNAR1 signaling in DCs resulted in a marked reduction in the number of T_H_2, TFH and CD44-hi CD4+ T cells in response to *Nb*. By contrast, cytokine responses to non-allergic stimuli such as DNFB, *Ms*, and *Ca* did not require IFNAR1 signaling, and Th2 responses to the contact sensitizer DBP-FITC were also independent of IFNAR1 signaling (12). Therefore, while IFN-I signaling in cDCs is not an essential requirement for TFH or Th2 differentiation (12) it can play an important role in some cases. This variable IFN-I requirement is likely due to differences in the skin cytokines that are elicited in each model (12, 44) and their impact on DC activation.

Previous publications have demonstrated a reduced IL-4 response to *Nb*, *Schistosoma mansoni* and HDM when IFNAR1 signaling was blocked (12, 13, 44). However, these experiments did not examine whether IFNAR1 signaling was facilitating the differentiation of GATA3-hi Th2 effector cells, or TFH which are major producers of IL-4 in LN (45), or both. Our experiments in aIFNAR1-treated mice and in IFNAR1^ΔCD11c^ vs. IFNAR^WT^ cKO mice (**Fig 1**) showed that GATA3-hi and total TFH populations both required IFNAR1 signaling. Experiments in *Il4*-AmCyan reporter mice showed that the differentiation of *Il4*-AmC+ TFH cells also required IFNAR1 signaling. IFNAR1 signaling in cDCs has been shown to be required for TFH differentiation after immunization with OVA+TLR ligands (28) and viral infection or vaccination (29), however, the role of IFN-I signaling in TFH differentiation to allergens has not been examined. We show that secondary *Nb* immunization of IFNAR1^ΔCD11c^ compared to IFNAR^WT^ mice led to a sustained defect in *Nb*-specific serum IgG1 and lower IgG1 affinity, thus extending previous findings to Th2 responses. Unlike specific IgG1, total serum IgE was similarly elevated in IFNAR1^WT^ and IFNAR1^ΔCD11c^ mice after secondary *Nb* immunization; further studies will be necessary to assess whether *Nb*-specific IgE is affected by IFN-I signaling in cDC2.

The reduction in the GATA3-hi and TFH CD4+ T cell response to *Nb* and HDM found in IFNAR1^ΔCD11c^ mice was due to a reduced number of divided cells *in vivo*, as shown by adoptive transfer of CTV-labelled polyclonal CD4+ T cells into IFNAR1^ΔCD11c^ or IFNAR^WT^ mice. Due to the small number of *Nb*-specific CD4+ T cells in the adoptively transferred population, it was not possible to establish whether this lower number of divided cells was due to insufficient stimulation and entry into cell division, or impaired survival of the divided cells (50). Due to the high proportion of cells undergoing 10+ divisions, estimates of fold-expansion and average number of divisions may well underestimate the differences between IFNAR1^ΔCD11c^ and IFNAR^WT^. While RT-qPCR experiments failed to demonstrate differential expression of the proliferative regulation genes *Myc* or *Bcl2* in fully divided CD4+ T cells, they did show decreased expression of genes involved in lactic acid and amino acid metabolism in IFNAR1^ΔCD11c^ compared to IFNAR^WT^ mice, suggesting a decreased use of cellular substrates for division (64). The increased expression of key T_H_2 and TFH genes in highly divided cells from IFNAR1^WT^ compared to IFNAR1^ΔCD11c^ mice (65) suggests additional defects in CD4+ T cell activation and/or differentiation that are independent of the lower number of divisions. The lower expression of *Il4ra* transcripts in the undivided population in IFNAR1^ΔCD11c^ hosts is also consistent with reduced IL-4-dependent signaling and lower IL-4 availability in these LNs (49).

### IFN-I-regulated genes explain the superior ability of IFN-I-conditioned cDC2s to induce T_H_2 differentiation

Despite the clear decrease in CD4+ T cell division observed in IFNAR^ΔCD11c^ mice compared to IFNAR^WT^, we found only moderate differences in the numbers of *Nb*-CTO+ cDC2s in LNs, and no significant differences in the expression of costimulatory markers and MHC-II on total or *Nb*+ migDC2 in LN by either flow cytometry or bulk RNAseq. These results are mostly consistent with our previous work showing similar expression of CD86 and PD-L2 on *Nb*+ migDC2 from aIFNAR-treated mice compared to isotype (12). Therefore, factors other than antigen availability and costimulation must be determining the lower response of IFNAR1^ΔCD11c^ vs IFNAR^WT^ mice to *Nb* immunization. Our analysis of global gene expression in *Nb*-CTO+ and PBS migDC2 from IFNAR^WT^ and IFNAR1^ΔCD11c^ mice showed the expected differences in expression of IFNAR-regulated genes, while transcripts for several cytokines and chemokines that have been associated with T_H_2 responses, such as *Ccl17* and *Ccl22*, and TFH responses eg *Il6*, were similarly expressed in IFNAR1^ΔCD11c^ and IFNAR^WT^ DC2, suggesting that other genes were responsible for the difference in T cell responses.

Amongst the DEGs that could explain the higher TFH and T_H_2 numbers in IFNAR1^WT^ vs IFNAR^ΔCD11c^ mice, *Arc* was upregulated in IFNAR1^WT^ compared to IFNAR1^ΔCD11c^ cDC2 and was demonstrated to promote rapid migration of cDCs to skin-draining LN (66), possibly explaining the lower numbers of *Nb+* DC in the IFNAR1^ΔCD11c^ skin-draining LN. *Il12b*, which was shown to inhibit T_H_2 activation *in vivo* (67, 68) was expressed at higher levels by CD11b-low cDC2 from IFNAR1^ΔCD11c^ mice compared to IFNAR1^WT^. It is however unclear whether this *Il12b* expression is sufficient to negatively regulate T_H_2 differentiation in IFNAR1^ΔCD11c^ mice. Transcripts for the cytokine *Il15* and its receptor *Il15ra* were also expressed at higher levels in IFNAR1^WT^ DC2 compared to IFNAR1^ΔCD11c^. IL-15 and IL-15Ra are upregulated following IFN-I signaling in cDCs (69) and increase T cell proliferation and survival (30). IL-15 was also reported to increase IL-4 and IFNγ expression by human (70) and mouse (63) CD4+ T cells activated *in vitro*, which may contribute to the increased T_H_2 responses observed in our experiments. Therefore, several genes that are differentially expressed in IFNAR1^WT^ compared to IFNAR1^ΔCD11c^ cDC2 could explain the superior ability of IFN-I-conditioned DC2 to induce T_H_2 and TFH differentiation.

*Ccl5* was also expressed at higher levels in CD11b-hi and CD11b-low cDC2 from *Nb*-immunized IFNAR1^WT^ mice compared to IFNAR1^ΔCD11c^. The receptor for CCL5, CCR5, is not expressed on T_H_2 or TFH cells but is found on many populations of immune cells including monocytes, and, together with direct IFN-I signaling (71, 72), might contribute to recruiting monocytes from tissues to IFNAR1^WT^ LNs (52). However, monocytes were found to be dispensable for robust GATA3-hi and IL-4 responses following intradermal *Nb* immunization (2), and were reported to suppress antibody responses to OVA (73) and LCMV (74). Thus, reduced monocyte numbers in LN are unlikely to explain the decreased T_H_2 and TFH responses in IFNAR1^ΔCD11c^ mice.

## Conclusions

This study provides a mechanism for the described role of IFNAR1 signaling in cDC2s in supporting T_H_2 immune responses after skin immunization with allergens including *Nb* and HDM, and identifies new candidate molecules that may play a role in IFN-I-dependent DC conditioning. IFN-I has emerged as an important modulator of the ability of cDC2s to prime T_H_2 responses in the LN, providing new insight into the T_H_2 priming capacity of skin cDC2s.

## Author Contributions

GRW, KLH and FR conceived the study and designed experiments; GRW, KLH, and SCT performed and analyzed *in vivo* and *in vitro* experiments; SIO analyzed and visualized bulk RNAseq data; OL analyzed RNAseq data and identified transcripts of interest; GRW and KLH drafted the manuscript and prepared the Figures; GRW, KLH, OL and FR edited the manuscript; all authors read and commented on the manuscript and approved the final draft for submission.

## Conflict of Interest Statement

The authors declare no conflicts of interest.

## Acknowledgements

The authors thank Samantha Small, Alix Grooby and Jonason Francisco of the Hugh Green Technology Centre (HGTC) for cell sorting, Alfonso Schmidt of the HGTC for aiding in BMDC imaging experiments, all the Biomedical Research Unit staff for mouse husbandry, and David O’Sullivan and Isabelle Montgomerie for helpful suggestions. The authors also thank Otago Genomics Ltd for advice on library preparation and sequencing of bulk RNA preparations.

This work was funded by a Health Research Council (HRC) of New Zealand Independent Research Organisation grant to the Malaghan Institute and an HRC project grant to FR. GRW and KLH were funded by PhD scholarships from the University of Otago Wellington, New Zealand. SIO was partly funded by the New Zealand Community Trust.

## Appendix

**Figure S1:**
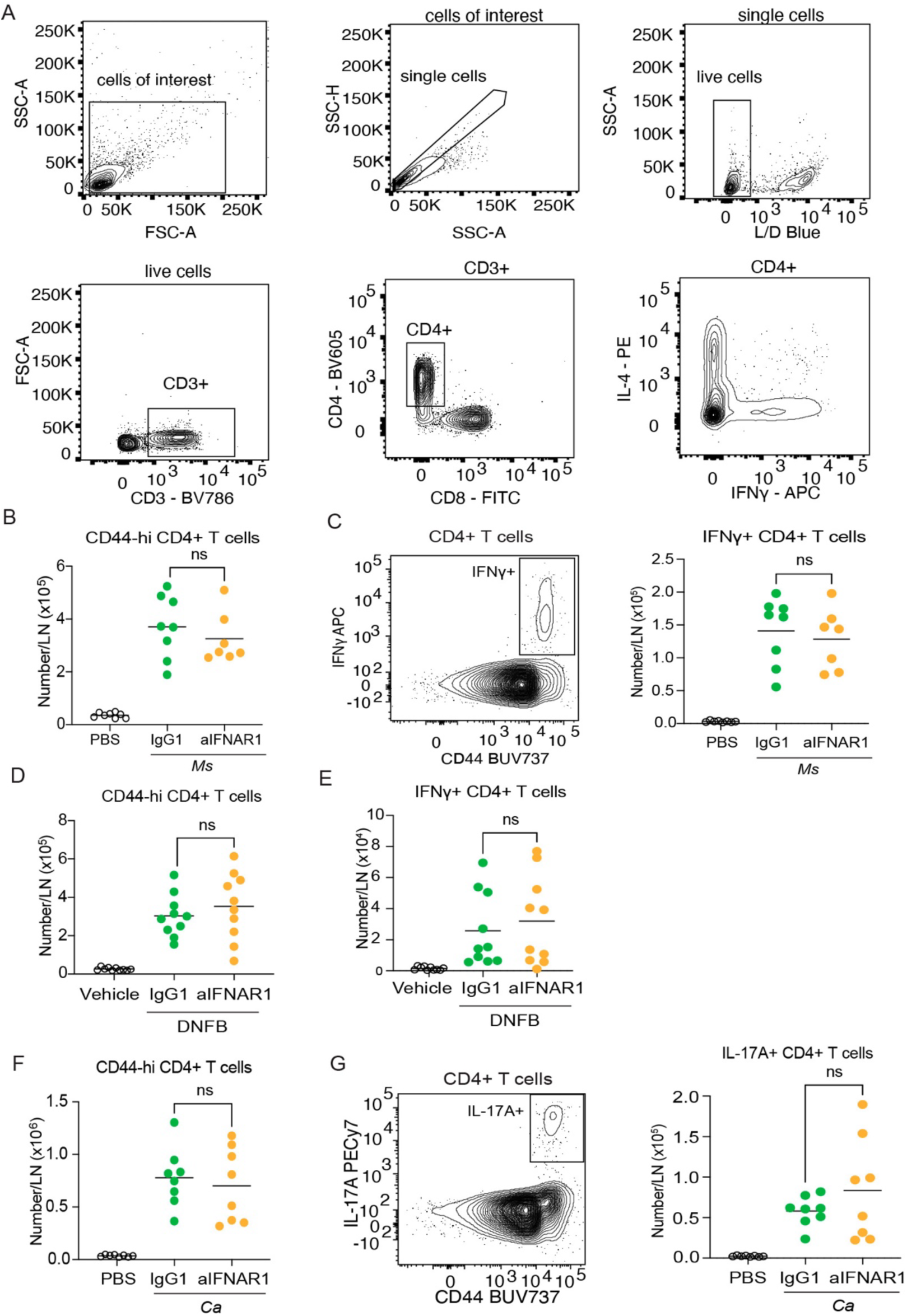
IFN-I signaling in DCs is not required for the development of IFNγ+ and IL-17A+ CD4+ cells. Mice were immunized intradermally with HDM, *Ms* and *Ca* or treated topically with DNFB. The ear-draining LN response was assessed by flow cytometry 7 days following immunization. **A:** Gating strategy for CD4+ T cells in the skin-draining LN. The lower right panel shows the relative lack of co-expression of IL-4 and IFNγ following immunization with HDM. **B:** Number of CD44-hi CD4+ T cells following *Ms* immunization. **C:** Representative gating and number of IFNγ+ CD4+ T cells following *Ms* immunization. **D:** Number of CD44-hi CD4+ T cells following DNFB treatment. **E:** Number of IFNγ+ CD4+ T cells following DNFB treatment. **F:** Number of CD44-hi CD4+ T cells following *Ca* immunization. **G:** Representative gating and number of IL-17A+ CD4+ T cells following *Ca* immunization. Dot plots show data from two pooled experiments each with 3 – 5 mice per group. Each dot corresponds to one mouse. *P* values refer to comparisons between the indicated groups and were calculated using the Mann-Whitney test. ns, *p*≥0.05.

**Figure S2:**
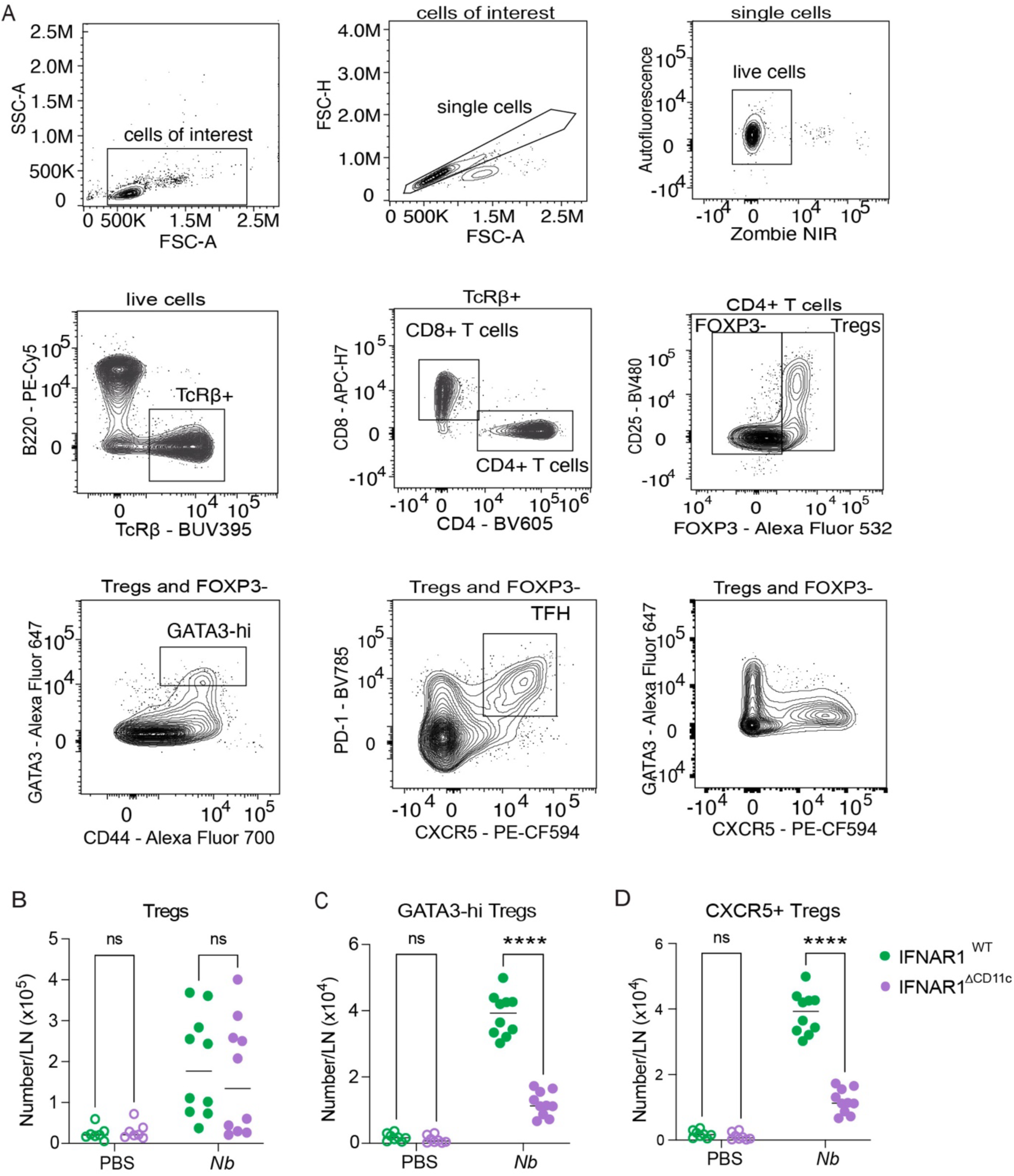
Regulatory T cell responses to *Nb* in IFNAR^WT^ and IFNAR^ΔCD11c^ mice. Mice were immunized intradermally with *Nb*, and the skin-draining LN response was assessed by flow cytometry 5 days following immunization. **A:** Gating strategy for CD4+ T cell responses in the skin-draining LN. Gating applies to **Fig 1F – H, 3B – D** and **S2B – E**. The lower right panel shows the relative lack of co-expression of CXCR5 and high levels of GATA3. **B:** Numbers of FOXP3+ Tregs. **C:** Numbers of GATA3-hi Tregs. **D:** Numbers of CXCR5+ Tregs. Dot plots show data from two pooled experiments each with 5 mice per group. Each dot corresponds to one mouse. *P* values refer to comparisons between the indicated groups and were calculated using two-way ANOVA. ****, *p*<0.0001; ***, *p*<0.001; **, *p*<0.01; *, *p*< 0.05; ns, *p*≥0.05.

**Figure S3:**
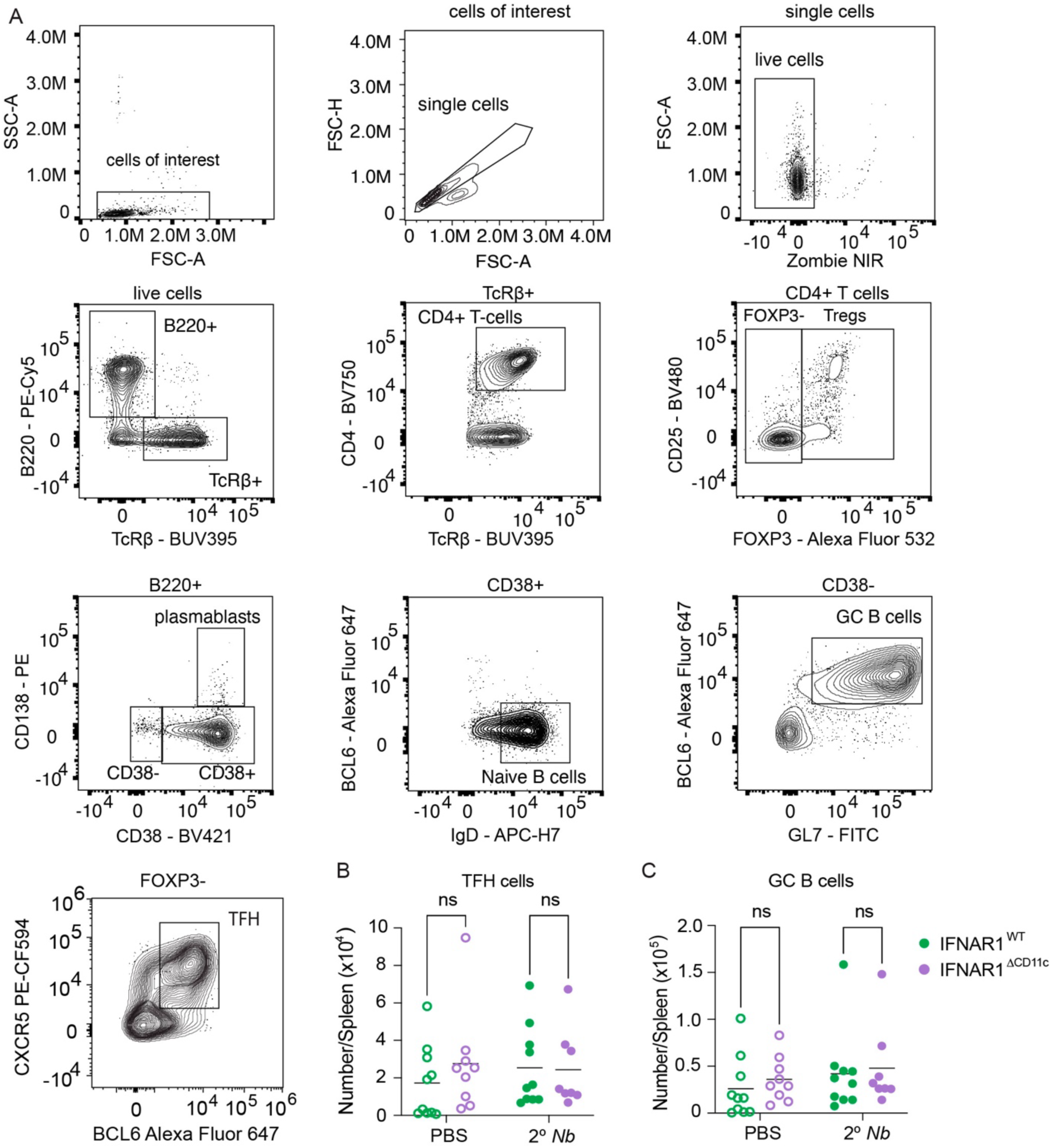
Lack of spleen germinal center responses after *Nb* intradermal immunization. IFNAR1^WT^ and IFNAR1^ΔCD11c^ mice were immunized intradermally with *Nb* on day 0 and day 21. The T and B cell response in the spleen was assessed on day 28. **A:** Gating strategy for the analysis of selected B cell and CD4+ T cell subsets in the skin-draining LN and spleen. Gating applies to **Fig 2D – E** and **S3B – C**. **B:** Numbers of spleen TFH. **C:** Numbers of spleen germinal center (GC) B cells. Dot plots show data from two pooled experiments each with 4 – 5 mice per group. Each dot corresponds to one mouse. *P* values refer to comparisons between the indicated groups and were calculated using two-way ANOVA. ns, *p*≥0.05.

**Figure S4:**
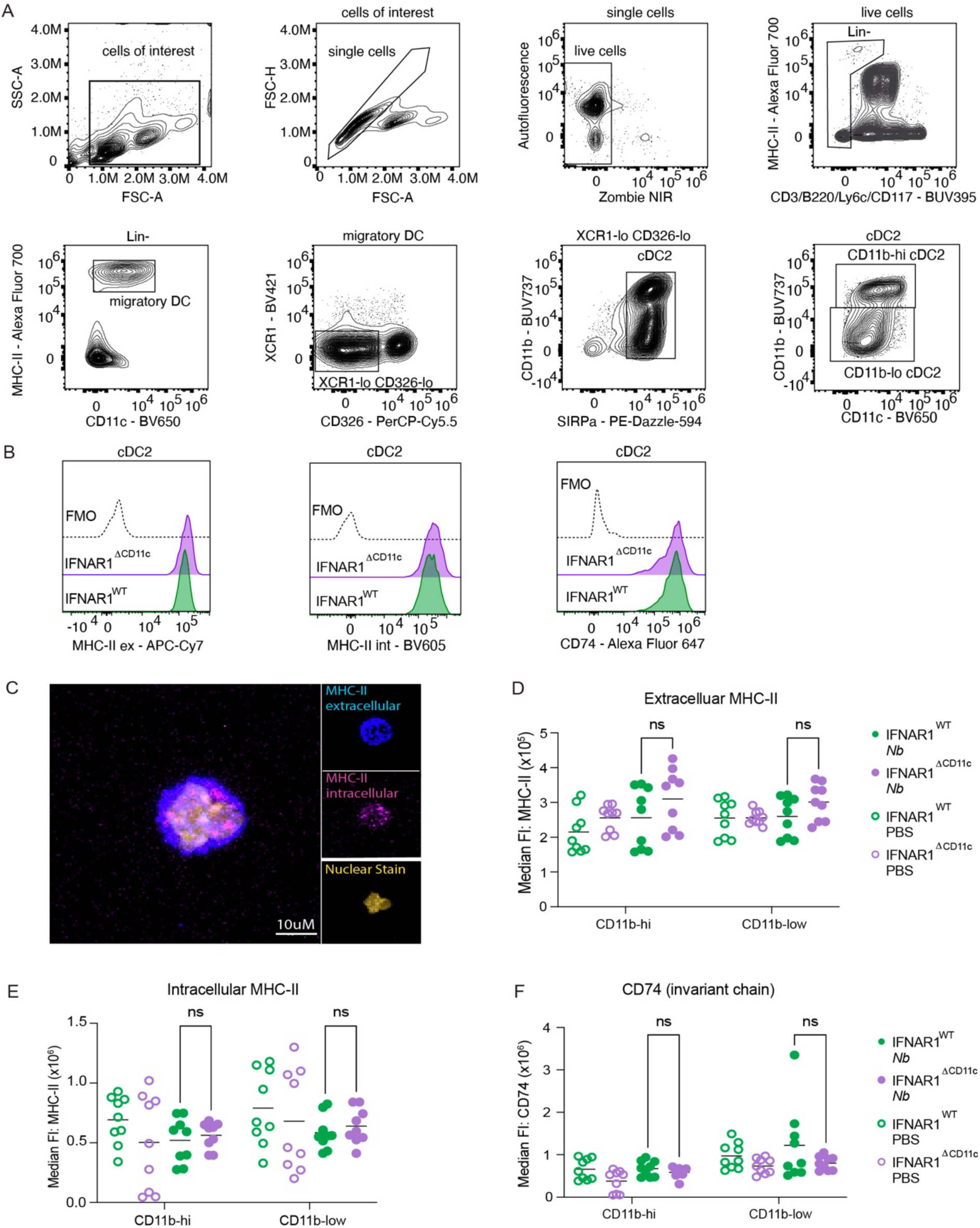
IFN-I signalling does not regulate MHC-II localization in cDC2s from *Nb*-immunized mice. **A:** Gating and cell sort strategy for the isolation of migratory cDC2 subsets from the skin-draining LN. Gating applies to **Fig 4A – F**, **S4C – E**, and **S5I** **B:** Representative histograms showing expression of extracellular and intracellular MHC-II and CD74 on skin-draining LN.migratory cDC2 subsets. **C:** Confocal microscopy of bone-marrow derived DCs to confirm extra-and intra-cellular staining of MHC-II molecules. Unstimulated BMDCs underwent a 10X expansion microscopy protocol and were stained for intra-and extracellular MHC-II. Nuclei were labelled using NuclearBlue prior to confocal microscopy. **D – F:** Mice were immunized intradermally with *Nb*, and the cDC2 response in the skin-draining LN was assessed two days following immunization. **D:** Median fluorescence intensity (MFI) of extracellular and **E:** intracellular MHC-II molecules on migratory cDC2s. **F:** MFI of CD74 on migratory cDC2s. Dot plots show data from two pooled experiments each with 4 – 5 mice per group. Each dot corresponds to one mouse. *P* values refer to comparisons between the indicated groups and were calculated using the multiple Mann-Whitney test. ns, *p*≥0.05.

**Figure S5:**
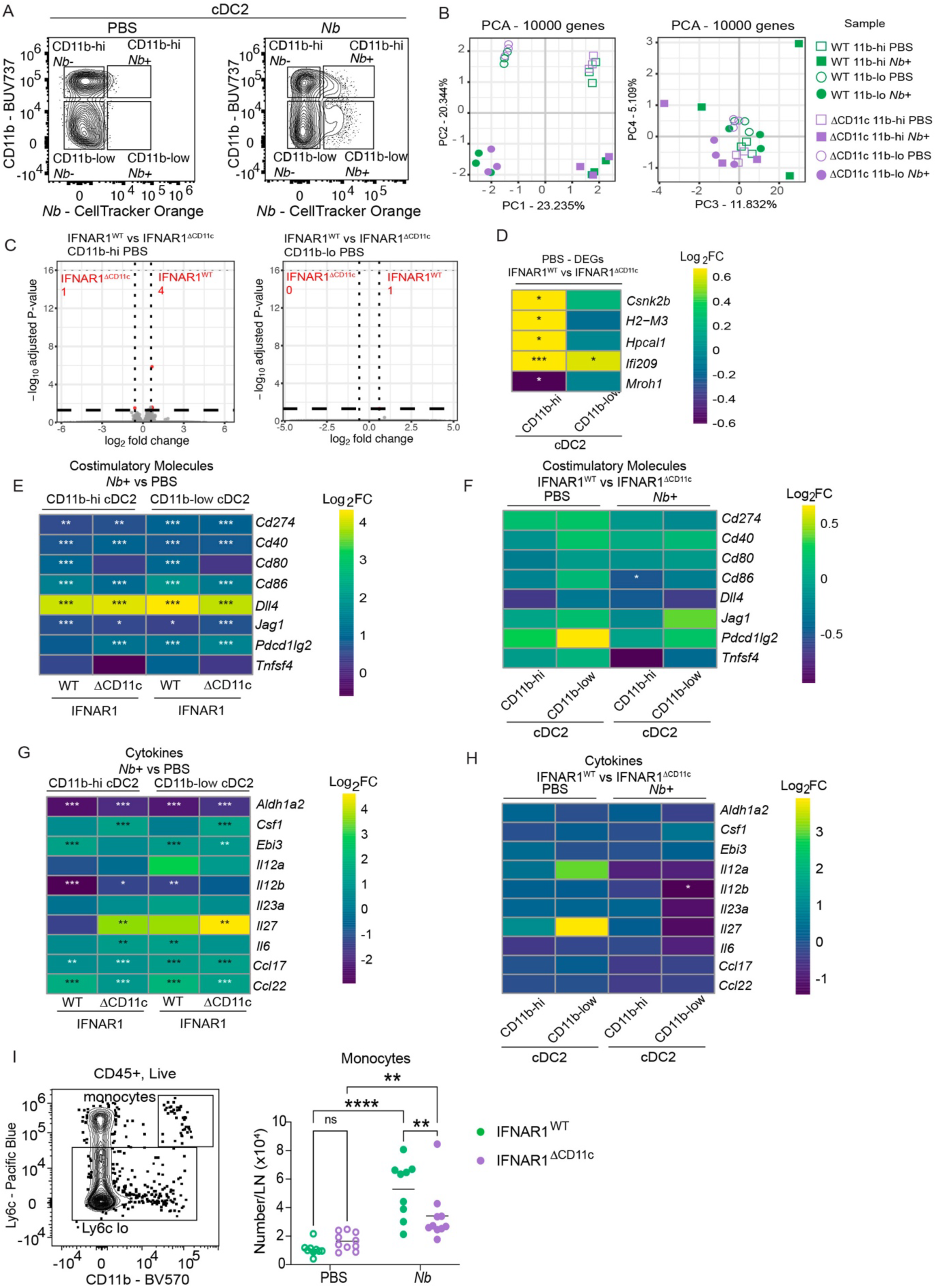
DEG and selected marker analysis of cDC2 subsets from *Nb*-immunized IFNAR1^WT^ and IFNAR1^ΔCD11c^ mice. IFNAR1^WT^ and IFNAR1^ΔCD11c^ mice were immunized intradermally with *Nb*-CTO, and *Nb*-CTO+ CD11b-hi and CD11b-low cDC2s were sorted and analysed by bulk RNA sequencing. **A:** Sorting strategy for Nb-CTO+ and Nb-CTO-CD11b-hi and CD11b-low cDC2s from IFNAR1^ΔCD11c^ mice **B:** Principal component analysis of the top 10,000 genes from Nb+ and PBS-treated CD11b-hi and CD11b-low migratory cDC2s from IFNAR1^WT^ and IFNAR1^ΔCD11c^ mice. **C:** Volcano plots showing DEGs in CD11b-hi and CD11b-low cDC2s from PBS-treated IFNAR1^WT^ vs IFNAR1^ΔCD11c^ mice. The numbers of DEGs are shown in the volcano plot. **D**: Heatmap of the DEGs in panel C. **E:** Heatmap of selected costimulatory marker expression in *Nb*+ vs PBS CD11b-hi or CD11b-low cDC2s from IFNAR1^WT^ and IFNAR1^ΔCD11c^ mice. **F:** Heatmap of selected costimulatory marker expression in IFNAR1^WT^ vs IFNAR1^ΔCD11c^ cDC2s that were either Nb-CTO+ or from PBS - treated mice. **G:** Heatmap of selected cytokine expression in *Nb*+ vs PBS CD11b-hi or CD11b-low cDC2s from IFNAR1^WT^ and IFNAR1^ΔCD11c^ mice. **H:** Heatmap of selected cytokine expression in IFNAR1^WT^ vs IFNAR1^ΔCD11c^ cDC2s that were either *Nb*-CTO+ or from PBS - treated mice. **I:** Representative gating and numbers of monocytes in the skin-draining LN of IFNAR1^WT^ vs IFNAR1^ΔCD11c^ mice two days following *Nb* immunization. The PCA (**B**) and heatmaps (**D - H**) show data from three biological replicates each corresponding to 2 – 3 pooled mice per group. *P* values in **D - H** refer to the indicated comparisons and were calculated using the DESeq2 Wald test. ***, *p*<0.001; **, *p*<0.01; *, *p*< 0.05. Dot plots (**I**) show data from two pooled experiments each with 4 - 5 mice per group. Each dot corresponds to one mouse. *P* values refer to comparisons between the indicated groups and were calculated using two-way ANOVA (**I)**. ****, *p*<0.0001; ***, *p*<0.001; **, *p*<0.01; *, *p*< 0.05; ns, *p*≥0.05.

## Notes

### Competing Interest Statement

The authors have declared no competing interest.

